# pH Induced Switch in the Conformational Ensemble of an Intrinsically Disordered Protein Prothymosin-*α* and Its Implications to Amyloid Fibril Formation

**DOI:** 10.1101/2022.06.25.497626

**Authors:** Lipika Baidya, Govardhan Reddy

## Abstract

Aggregation of intrinsically disordered proteins (IDPs) is the cause of various neurodegenerative diseases. Changes in solution pH can trigger IDP aggregation due to a shift in the IDP monomer population with a high aggregation propensity. Although there is experimental evidence that acidic pH promotes the compaction of IDP monomers, which subsequently leads to aggregation, the general mechanism is not clear. Using the IDP prothymosin-*α* (proT*α*), which is involved in multiple essential functions as a model system, we studied the pH effect on the conformational ensemble of proT*α* and probed its role in aggregation using a coarse-grained IDP model and molecular dynamics simulations. We show that compaction in the proT*α* dimension at low pH is due to the protein’s collapse in the intermediate region (E41 - D80) rich in glutamic acid residues. Further, the *β*-sheet content increases in this region upon pH change from neutral to acidic. We hypothesized that the conformations with high *β*-sheet content could act as aggregation-prone (*N*^*∗*^) states and nucleate the aggregation process. We validated our hypothesis by performing dimer simulations starting from *N*^*∗*^ and non-*N*^*∗*^ states. We show that simulations initiated using *N*^*∗*^ states as initial conformations form dimers within 1.5 *μ*s, whereas the non-*N*^*∗*^ states do not form dimers within this timescale. This study contributes to understanding the general principles of pH-induced IDP aggregation. The main result upon pH change from neutral to acidic, the intermediate region of proT*α* is responsible for aggregation due to an increase in its *β*-sheet forming propensity and forms the fibril core can be verified by experiments.

**Graphical TOC Entry:** 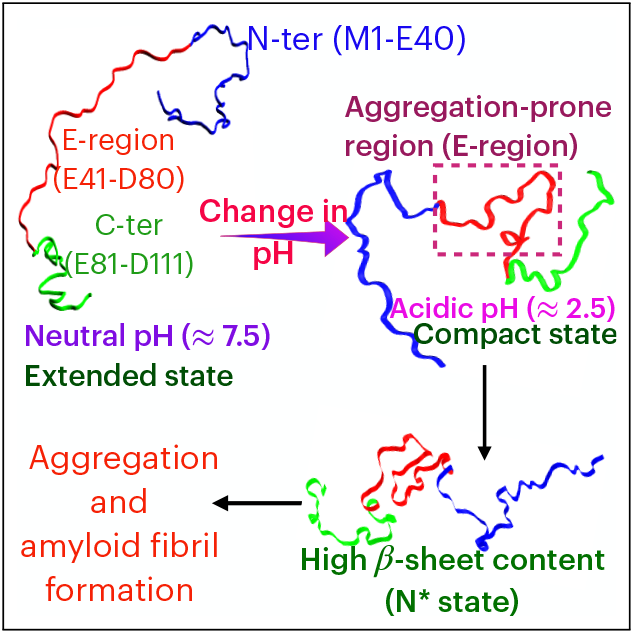

---

Intrinsically disordered proteins (IDPs)^1–3^ are a class of proteins that lack secondary and tertiary structures and exist as an ensemble of inter-converting conformations. Despite being disordered, IDPs are involved in various biological functions^4–6^ such as cell regulation, signal transduction, transcription etc. Aberrant behavior of IDPs forming amyloid fibrils through aggregation leads to diseases^7,8^ such as Alzheimer’s, Parkinson’s, tauopathy, and cancer. External factors^9^ such as temperature, ^10^ pH,^11,12^ salts, ^13,14^ cosolvents, ^15–17^ crowder^17,18^ etc. modulate the conformational ensemble as well as the aggregation propensities ^18–23^ of IDPs.

Prothymosin-*α* (proT*α*)^24^ is a 111-residue acidic nuclear IDP involved in various functions^25,26^ such as cell proliferation, chromatin remodeling and proapoptotic activity. Over-expression of proT*α* in cells are linked to pulmonary emphysema, cancer, polycystic kidney disease, tumor aggressiveness and poor prognosis. Circular dichroism (CD) experiments^11,24^ showed that proT*α* mostly exists as a random coil due to a high fraction of negatively charged residues. The extended and disordered conformations of proT*α* are responsible for its various functions. ^25,26^ The conformational ensemble of proT*α* is modulated using salt and pH through the variation of electrostatic interactions. ^11,13,14^ Small-angle X-ray scattering (SAXS) experiments^11^ showed that proT*α* changes its conformation from random coil to a partially collapsed state on pH change from neutral to acidic. It was also observed that at low pH, proT*α* aggregates into regular elongated amyloid fibrils. ^27^ However, the relation between the partially collapsed state and the aggregation of proT*α* into fibrils at acidic pH is elusive.

Previous studies^28–34^ on both globular proteins and IDPs showed that the aggregation propensities are encoded in the monomer conformational ensemble. It was further demonstrated that the aggregation-prone monomer states (*N*^*∗*^) have similar characteristics to the fibril state and are masked in the monomer conformational ensemble. ^29,31–35^ Simulations^30,33^ using lattice models and all-atom description demonstrated that the timescale for fibril formation strongly depends on the population of *N*^*∗*^ states. Therefore, it is essential to elucidate the *N*^*∗*^ states to decode the pH-dependent protein aggregation rate. We asked the following questions to understand the pH-dependent proT*α* aggregation: 1) What part of the proT*α* contributes to the compaction upon pH change from neutral to acidic? 2) How does the propensity of the secondary structure formation change with pH and its role in inducing aggregation? 3) What are the characteristics of aggregation-prone (*N*^*∗*^) conformations? 4) How does the *N*^*∗*^ state influence the aggregation process?

Experiments proposed that the partially collapsed states initiate the aggregation of proT*α*.^11,27^ However, it is difficult to elucidate the structural details of aggregation-prone states from experiments due to the transient nature of IDPs. Computer simulations can complement experiments to understand the IDP conformational ensemble ^36–43^ and probe the aggregation^15,44–47^ mechanism. In this work, we used a coarse-grained two bead model, ^14,36^ known as the self-organized polymer model for IDP (SOP-IDP), to study the pH effect on the conformational ensemble of proT*α* and its aggregation. The change in pH from neutral (pH ≈ 7) to acidic (pH ≈ 2) is taken into account by varying the charged states of negatively charged residues. On changing pH from neutral to acidic, we observed that the conformation ensemble of proT*α* undergoes compaction. Compaction, as well as *β*-sheet content at low pH, is maximum in the intermediate region (E41-D80) of proT*α* rich in glutamic acid residues compared to the N-terminal (M1-E40) and C-terminal (E81-D111). We showed that the conformations with high *β*-sheet propensities act as the *N*^*∗*^ states. We provide evidence that the *N*^*∗*^ states form dimers faster than the non-*N*^*∗*^ states and can be critical to nucleation in the early stages of aggregation.

## Kratky plots of proTα at neutral and acidic pH

ProT*α* is a charged IDP of length, *N*_*res*_ = 111 residues and is dominated by negatively charged residues. The fraction of positively charged residues (*f*_+_) and negatively charged residues (*f*_−_) in proT*α* are 0.09 and 0.49, respectively. ProT*α* can be classified as a strong polyelectrolyte^48,49^ as the fraction of charged residues (*FCR* = *f*_+_ + *f*_−_ = 0.58) is greater than 0.3, and the net charge per residue^48–50^ (*NCPR* = |*f*_+_ − *f*_−_| = 0.4) is greater than 0.

To study the pH effect on the conformational ensemble of proT*α*, we performed Langevin dynamics simulations at temperature *T* = 298 K, and monovalent salt of concentration, [*salt*] = 150 mM. Experiments typically use monovalent salt in the concentration range, [*salt*] = 100 - 200 mM.^11,24^ We changed the charged state of the negatively charged residues to neutral to mimic the pH change in experiments^11^ from 7.5 to 2.5. Accounting for the pH change by varying the charge is a reasonable assumption in this case as the acidic residues are dominantly in the protonated state at pH ≈ 2.5. The p*K*_a_ of aspartic acid (Asp) and glutamic acid (Glu) side chains are 3.9 and 4.3, respectively. The fraction of Asp and Glu sidechains in the deprotonated state (*f*_*Dprot*_) can be calculated using the Henderson-Hasselbalch equation,

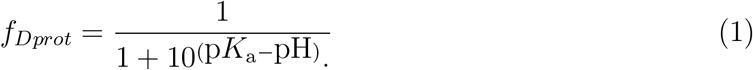

The fraction of Asp and Glu side chains in the protonated state (1-*f*_*Dprot*_) at pH ≈ 2.5 are 96.17 % and 98.44 %, respectively, and their charge is taken to be 0. At pH ≈ 7.5, the side chains of Glu and Asp are dominantly in the deprotonated state, and their charge is -1. The charged state of positively charged residues is +1 at both neutral and low pH.

We computed the normalized Kratky plot, *q*^2^*I*(*q*)*/I*(0) as a function of *q*, where *I*(*q*) and *I*(0) are the scattering intensities at wave vectors *q* and *q* = 0, respectively (Figure 1A). The near quantitative agreement between the computed and experimental Kratky plots shows that the assumptions we made are reasonable and the SOP-IDP model captures the changes in the proT*α* conformational ensembles with the pH change from neutral to acidic.

**Figure 1:**
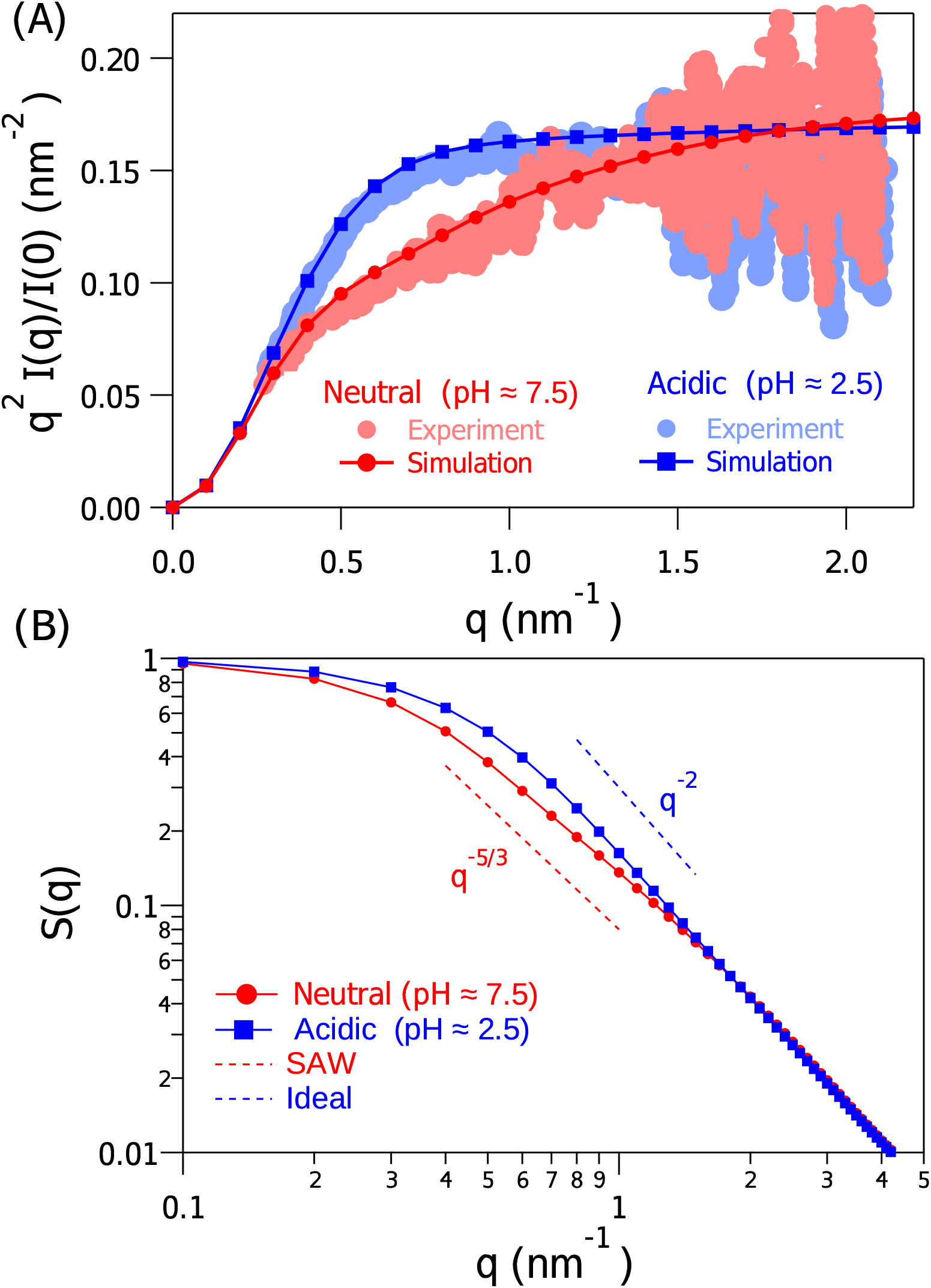
(A) Normalized Kratky plot (*q*^2^*I*(*q*)*/I*(0) vs *q*) from experiments (red and blue circles) and simulations (solid line with red circles and blue squares) for proT*α* at *T* = 298 K, monovalent [*salt*] = 150 mM, and pH ≈ 7.5 (red) and 2.5 (blue). (B) Simulated structure factor *S*(*q*) for proT*α* at pH ≈ 7.5 (solid line with red circles) and 2.5 (solid line with blue squares). The dashed straight lines with scaling *q*^−5*/*3^ and *q*^−2^ correspond to the self-avoiding walk (SAW) (red) and ideal chain (blue) configuration.

At neutral pH, the Kratky plot increased monotonically with *q*, which is typical for a disordered IDP (Figure 1A). However, at low pH, the Kratky plot increased faster compared to neutral pH in the low *q* region (0 nm^−1^ < *q* < 0.7 nm^−1^), and reaches a plateau at high *q* region (*q* > 1.2 nm^−1^) (Figure 1A). The change in the Kratky plot from a monotonically increasing plot at neutral pH to a relatively flat curve for larger *q* at lower pH suggests that there is a transition in the conformational ensemble from disordered/extended state to a relatively compact state^11^ (Figure 1A). We also assessed the pH effect on the conformational ensemble of proT*α* by extracting the Flory scaling exponent, *ν* from the structure factor ^51^ *S*(*q*) (Eq. 4) using the relation, *S*(*q*) ∼ *q*^−1/*ν*^. The scaling exponent varied from *ν* ≈ 0.6 to *ν* ≈ 0.5 with pH change from neutral to acidic (Figure 1B). At neutral pH, although the net charge on proT*α* is -44, it behaves as a self-avoiding walk (SAW) instead of a rod-like chain due to the screening of electrostatic interactions by the salt ([*salt*] = 150 mM). In the absence of the salt at neutral pH, proT*α* behaves as a rod-like chain with *ν* ≈ 1.0.^14^ The CD and SAXS^11,24^ experiments also confirm that proT*α* exists in an extended state at neutral pH. However, the negatively charged residues become protonated with decreasing pH, and the chain size compacts to behave like an ideal chain (Figure 1B). In simulations, the average radius of gyration (⟨*R*_*g*_⟩) of the IDP decreased from 38.3 Å to 30.7 Å on changing pH from neutral to acidic, which is in near quantitative agreement with the SAXS experiments.^11^ In the SAXS experiments,^11^ on changing pH from 7.5 to 2.5, *R*_*g*_ decreased from ≈ 38 Å to ≈ 28 Å.

## ProTα compacts in the E-Region (E41 - D80) at low pH

The conformational ensemble of proT*α* is highly heterogeneous^36,52–55^ due to its heteropolymeric nature. Therefore, global properties such as the Kratky plot or ⟨*R*_*g*_⟩; are insufficient to provide the microscopic picture of the varying conformational ensemble with changing pH. We computed the difference in the probability of contact formation among all the residues at neutral and acidic pH to assess the pH effect on proT*α* conformations (Figure 2A). If the distance between the two residues is less than a cutoff distance (≤ 8 Å), we consider those residues to be in contact. Depending on the change in contact frequency, we categorized the full-length proT*α* into three segments - (1) residues M1 to E40 denoted as N-terminal (N-ter), (2) residues E41 to D80 denoted as E-region (due to the presence of a high fraction of glutamic acid), and (3) residues E81 to D111 denoted as C-terminal (C-ter). With decreasing pH, we observed that the contact frequency decreased in the two terminal regions and increased in the E-region (Figure 2A).

**Figure 2:**
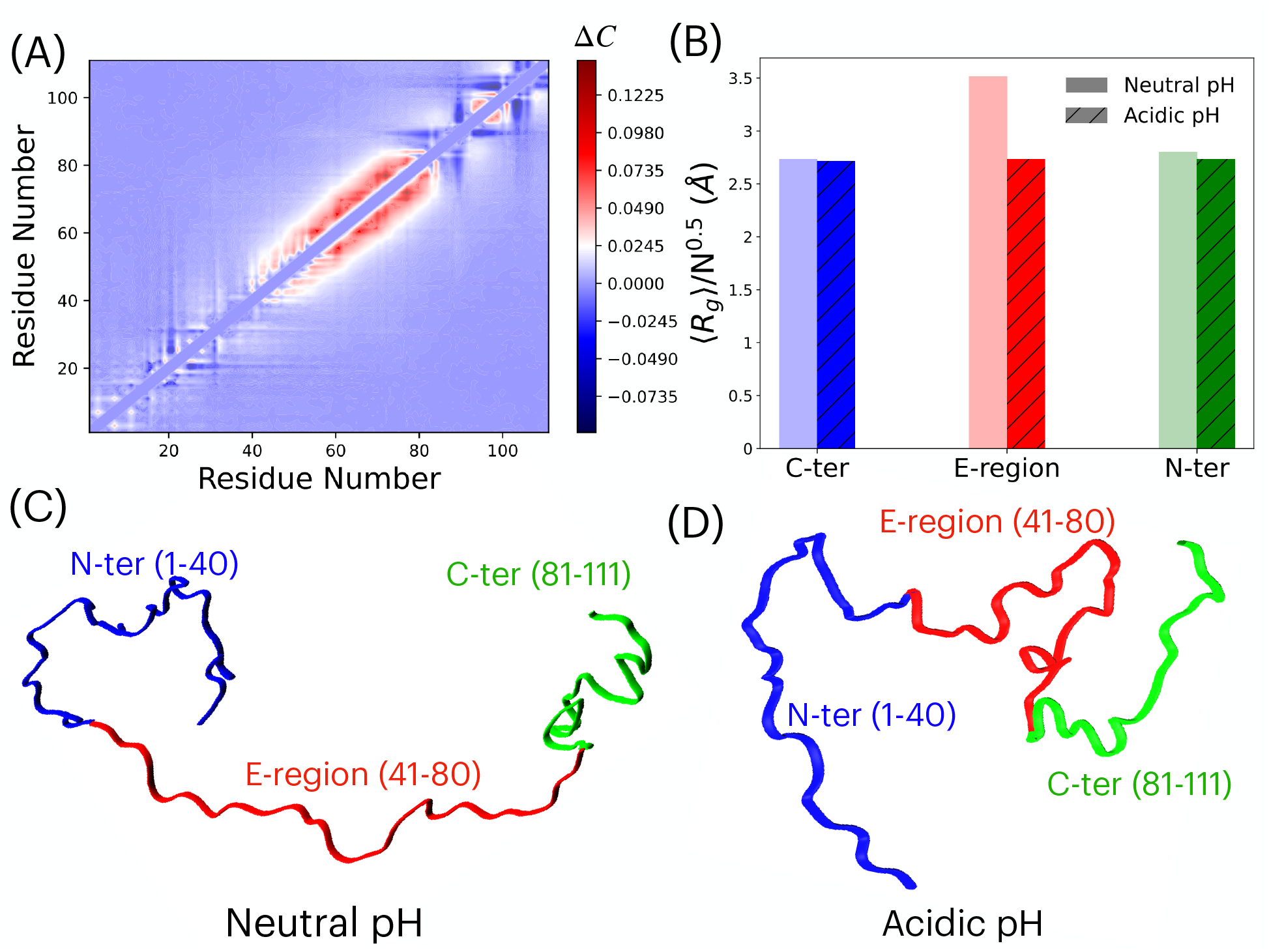
Effect of pH on the three segments of proT*α*. (A) Difference in proT*α* contact map 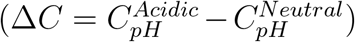 obtained at acidic pH (pH ≈ 2.5) and neutral pH (pH ≈ 7.5). The color bar denotes the change in the probability of contact formation (Δ*C*). Positive values (red) correspond to an increase in the frequency of contact formation, and negative values (blue) correspond to a decrease in the frequency of contact formation upon pH change. (B) The rescaled radius of gyration, ⟨ *R*_*g*_ */N* ^0.5^ ⟩, where *N* is the segment length, is plotted for neutral and acidic pH for all the three segments. The light shaded bar and solid bar with line correspond to neutral and acidic pH, respectively. To minim<u>iz</u>e the effect of segment length in comparing the ⟨ *R*_*g*_ ⟩ changes, we rescaled the ⟨ *R*_*g*_ ⟩ with 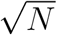, accounting for an ideal chain of length *N*. Representative conformations of proT*α* at neutral and acidic pH are shown in (C) and (D), where the three segments are denoted as N-ter (M1 - E40) in blue, E-region (E41 - D80) in red, and C-ter (E81 - D111) in green.

At neutral pH, the C-ter and N-ter region of the protein is composed of both positively and negatively charged residues. In contrast, the E-region is enriched with negatively charged residues, especially glutamic acid (E). On reducing pH, the electrostatic attraction between the oppositely charged residues present in both N-ter and C-ter vanishes, and the repulsion among the positively charged residues increases, which leads to a lowering in contact frequency (Figure 2A,C,D). In contrast, there is a strong repulsion between the negatively charged residues present in the E-region at neutral pH, which diminishes at low pH due to charge neutralization^56^ (Figure 2A,C,D).

We quantified the effects of pH change on the three segments of proT*α* by computing the relative change in ⟨*R*_*g*_⟩. On changing pH from neutral to acidic, the decrease in ⟨*R*_*g*_⟩ for N-ter and C-ter are ≈ 0.2 Å and 0.4 Å, respectively which is insignificant. However, lowering pH led to a significant compaction in the E-region, and ⟨*R*_*g*_⟩ decreased by ≈ 5 Å (Figure 2B). The extent of compaction for each segment is governed by its NCPR. ^57,58^ At neutral pH, the NCPR for N-ter, E-region, and C-ter are 0.15, 0.7 and 0.32, respectively, and these values at low pH decrease to 0.12, 0, and 0.16. The change in contact frequency and extent of compaction for each segment is directly proportional to the change in NCPR values (0.03, 0.7, and 0.16 for N-ter, E-region, and C-ter, respectively) due to the switch in pH conditions from neutral to acidic. ^56–58^ We conclude that lowering pH modulates the contact probability^59–61^ significantly in the E-region and hypothesize that this sequence with altered charges is probably conducive to secondary structure formation.

## E-Region is conducive for secondary structure formation at low pH

We quantified the structure formation with varying pH from simulations using the program PCASSO. ^62^ PCASSO is a fast and accurate machine learning-based method to assign secondary structural elements (SSE) such as helix, *β*-strand, and coil for each residue in the protein chain using only the *C*_*α*_ atom coordinates. The method achieves ≈ 95 % accuracy compared to the commonly used SSE calculation algorithm DSSP. ^63^ We computed the percentage of SSE formation (helix/*β*-strand/coil) for each residue of proT*α* using the conformations obtained from simulation trajectories and the PCASSO program. Most of the residues prefer coil state (> 97 %) at both neutral and low pH (Figure S1A). However, the percentage of helix and *β*-sheet formation are not negligible (< 2 %) (Figure 3 and S1). We observed that lowering pH increased the amount of helix or *β*-sheet content (Figure 3 and S1B). Our observation is in qualitative agreement with the experiments, ^11,27^ which reported ≈ 10 % SSE at low pH.

**Figure 3:**
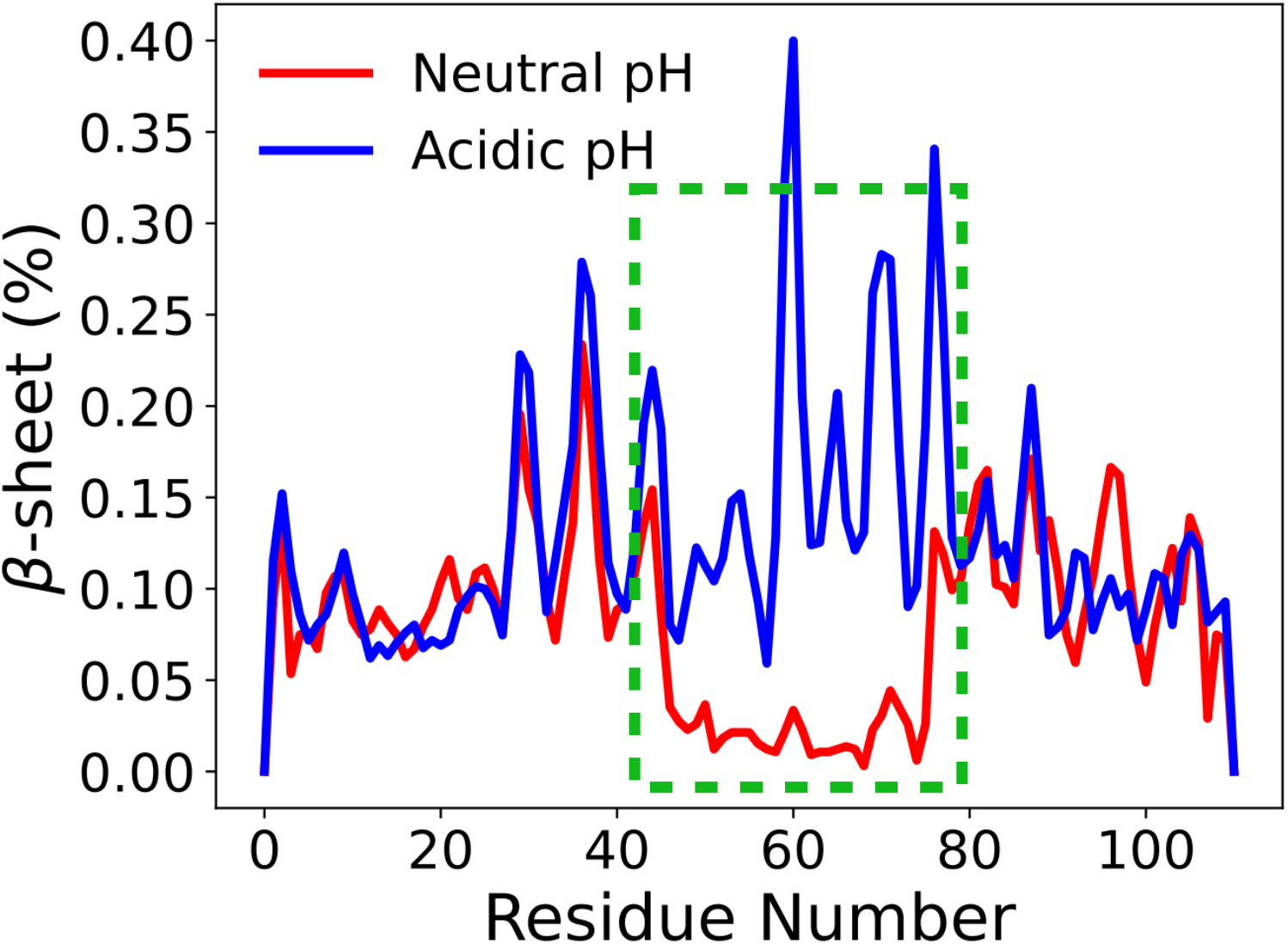
The percentage of *β*-sheet content is plotted as a function of proT*α* residues at neutral pH (red) and acidic pH (blue). The *β*-sheet content increased at acidic pH in the E-region (E41 - D80) highlighted using a green rectangle.

The *β*-sheet and helix content in C-ter and N-ter regions vary between 0.05 - 0.3 % and 1 - 2 % at neutral pH, respectively. At acidic pH (≈ 2.5), the SSE content remains almost similar to neutral pH in both regions (Figure 3). However, the E-region has a noticeable increment in SSE on pH change (Figure 3). At neutral pH, *β*-sheet/helix content in E-region is ≈ 0. The E-region exists as a coil due to the electrostatic repulsion between the charged glutamic acid residues. With the decrease in pH, the charges on glutamic acid are neutralized, and the propensity for *β*-sheet formation is enhanced. For some residues in the E-region (E59-G60, E69-G70, and D76-G77), the *β*-sheet content is 2-3 times higher than C-ter and N-ter (Figure 3). Enhancement in *β*-sheet content in E-region suggests that the effect of pH switch from neutral to acidic is maximum for this region and can initiate aggregation.

## Ensemble of N ^∗^ state conformations is heterogeneous

The simulations in this study show that at acidic pH, proT*α* compacts in the E-region, and enhanced *β*-sheet content is observed in this region. Earlier studies have shown that an increase in *β*-sheet content is directly proportional to the aggregation propensity of proteins.^47,64^ We hypothesized that enhanced propensity for *β*-sheet formation in the E-region leads to proT*α* aggregation, and E-region acts as a fibril forming core. The conformations with enhanced *β*-sheet content in the E-region can act as an aggregation-prone *N*^*∗*^ state^28,29^ and initiate the aggregation process.

We used each residue’s *β*-sheet content information to assign whether a conformation belongs to the aggregation-prone *N*^*∗*^ state.^65^ If a single residue present in the E-region of a conformation obtained from the simulations is assigned to a *β*-sheet by PCASSO then we designated that conformation belongs to the *N*^*∗*^ state. Previous studies ^29–35^ on both globular proteins and IDPs have demonstrated that *N*^*∗*^ states are present within the monomer ensemble and the population of *N*^*∗*^ states dictates the aggregation rate.

We characterized the *N*^*∗*^ conformations using t-distributed stochastic neighbor embedding (t-SNE) non-linear dimensionality reduction technique,^66^ which reduces dimensionality while maintaining the relationship between the data points. We used the pairwise distance between backbone beads to project high dimensional data into low dimensional space. The dimensionality reduction analysis revealed two distinct clusters. The population of cluster-1 is ≈ 61 % and cluster-2 is ≈ 39 % (Figure 4). We projected *R*_*g*_ on each data point to identify the size of the *N*^*∗*^ conformations in each cluster. The *R*_*g*_ shows that cluster-1 and cluster-2 are dominated by compact and extended conformations, respectively, and suggests that the *N*^*∗*^ ensemble is comprised of both compact and extended conformations. However, the dominating population of *N*^*∗*^ states is compact (≈ 61 %), and it agrees with the experiment, which inferred that compact states mostly drive the aggregation process.^15,27^ The probability distribution of *R*_*g*_, *P* (*R*_*g*_), shows that the range of *R*_*g*_ values for conformations within cluster-1 and cluster-2 are between 18 - 30 Å and 30 - 52 Å, respectively (Figure S2). The *P* (*R*_*g*_) shows that both the clusters contain a range of conformations with various sizes indicating that the *N*^*∗*^ ensemble is heterogeneous in nature.

**Figure 4:**
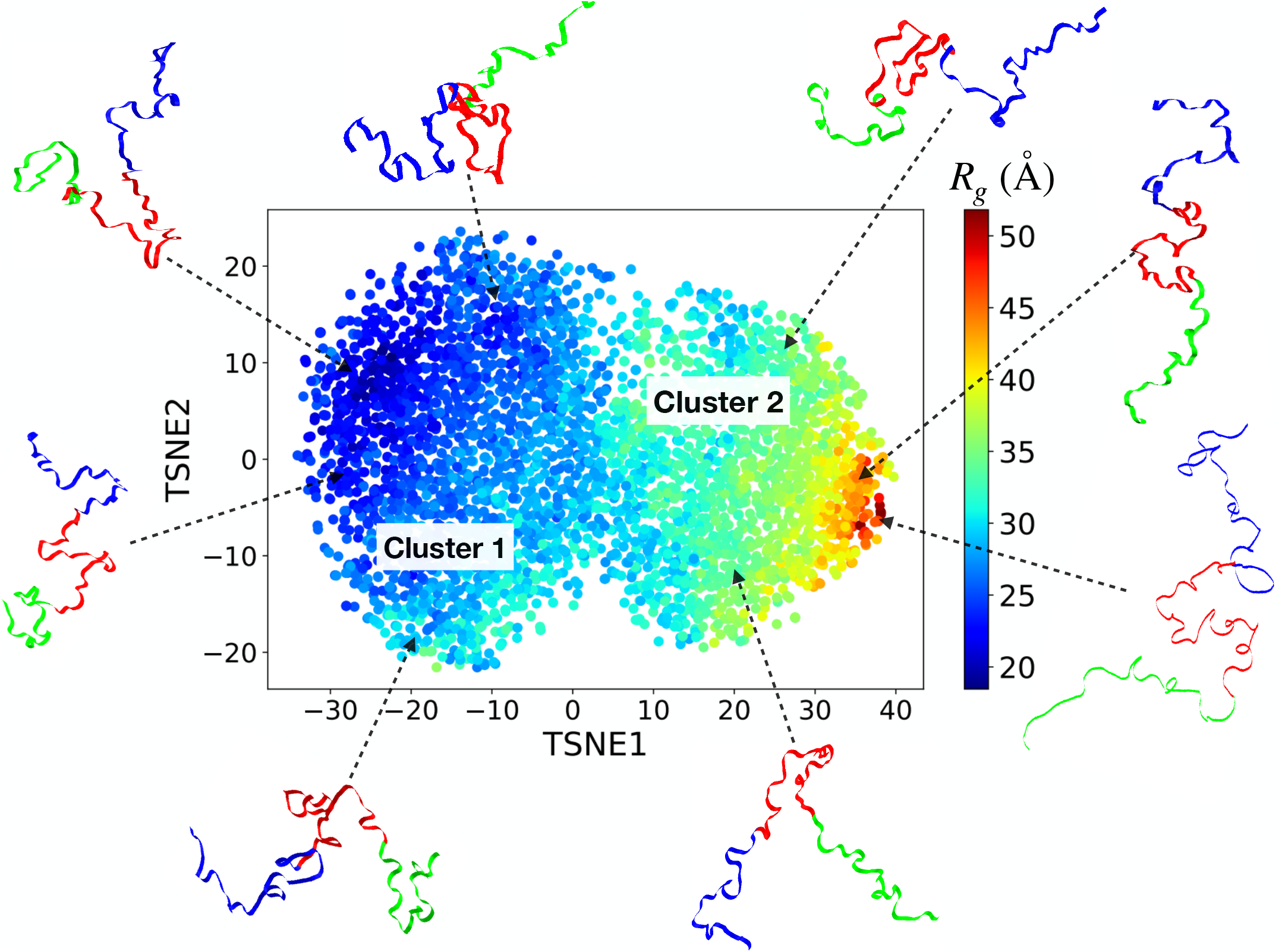
t-SNE dimensionality reduction analysis of the *N*^*∗*^ state conformations. Two distinct clusters are dominated by the compact (cluster 1) and extended (cluster 2) conformations. The color of each data point shows the *R*_*g*_ value corresponding to the conformation of that data point. The representative *N*^*∗*^ structures of proT*α* are shown in ribbon representation, where the N-ter, E-region, and C-ter are shown in blue, red, and green, respectively.

## N ^∗^ conformations readily form dimers at low pH

The amyloid fibrils of proT*α* obtained at low pH *in vitro* are characterized using scanning force and electron microscopy.^27^ Experiments showed that proT*α* amyloid fibrils completely break down for pH > 5. The proposed mechanism for fibril formation is that at low pH, there is a shift in the proT*α* monomer population from random coil to partially compact states, which can form *β*-strand rich nuclei to initiate oligomer formation. ^27^

We probed the aggregation propensity of *N*^*∗*^ states by performing dimer simulations. We randomly chose two conformations and initiated the simulation by placing them at *R*_*g*_ distance (≈ 30 Å) to maintain the critical concentration for dimer formation. To track the dimerization process, we calculated the fraction of inter-chain contact formation 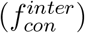 between two chains in the E-region. If the distance between either the backbone or the side chain bead of a residue belonging to chain-1 is within 8 Å of the backbone or the side chain bead of a residue belonging to chain-2, then we consider that those two residues are in contact and denote it as an interchain contact. There are 41 residues in the E-region of proT*α*, and the maximum number of interchain residue contacts possible is *N*_*max*_ ≈ 41, assuming that one residue is in contact with only one another residue. Although with a 8 Å cutoff distance it is possible that a residue can be in contact with multiple other residues. However, we counted such residues only once while computing the number of interchain residue contacts. We considered the proteins to be in the dimer state if the number of interchain residue contacts (*N*_*inter*_) exceeded 50 % of *N*_*max*_ during the dimer simulation. We defined the fraction of interchain contacts for a conformation using the relation, 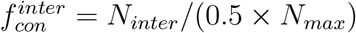. The dimer state corresponds to 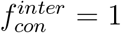, and the monomer state corresponds to 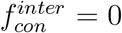. We initiated 50 independent simulations for each set of dimerization conditions studied.

We compared the dimer formation propensity of proT*α* at both neutral and acidic pH by plotting 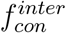 as a function of the simulation time. At neutral pH, 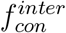 fluctuates between 0 to 0.4 and does not approach 1, suggesting that there are not enough interchain interactions between the two chains to form the dimer (Figure S3). We did not observe any dimer formation event for any trajectories at neutral pH. This observation agrees with experiments, ^27^ which show that proT*α* does not aggregate at neutral pH.

To validate our hypothesis that *N*^*∗*^ states with *β*-sheet content are responsible for aggregation, we initiated dimer simulations starting with *N*^*∗*^ as well as non-*N*^*∗*^ states at acidic pH. We tracked the dimer formation events by computing 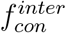 as a function of time. When we initiated the simulations using *N*^*∗*^ conformations, the interchain contacts between the two chains formed and ruptured multiple times due to the dynamic nature of dimer formation and the absence of multiple protein chains. The 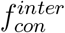 mostly oscillates between 0.4 - 1, ensuring that *N*^*∗*^ states form dimer spontaneously on the timescale of microseconds and mostly remain in a dimer state. Dimerization was observed for ≈ 68 % trajectories starting with *N*^*∗*^ states within 4.5 *μ*s and in majority of the simulations dimers form 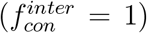 within 0.12 - 1.5 *μ*s (Figure 5A). However, in simulations initiated using the non-*N*^*∗*^ states at acidic pH, the 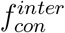 does not approach 1 and fluctuates between 0 - 0.4, which shows that non-*N*^*∗*^ states are not efficient in dimer formation and initiating aggregation. We did not observe the dimerization event in ≈ 87 % trajectories initiated with non-*N*^*∗*^ states on the same 4.5 *μ*s time scale (Figure 5). The frequency of contact formation for non-*N*^*∗*^ state at acidic pH is higher than neutral pH but not enough to form dimers (Figure 5 and S3).

**Figure 5:**
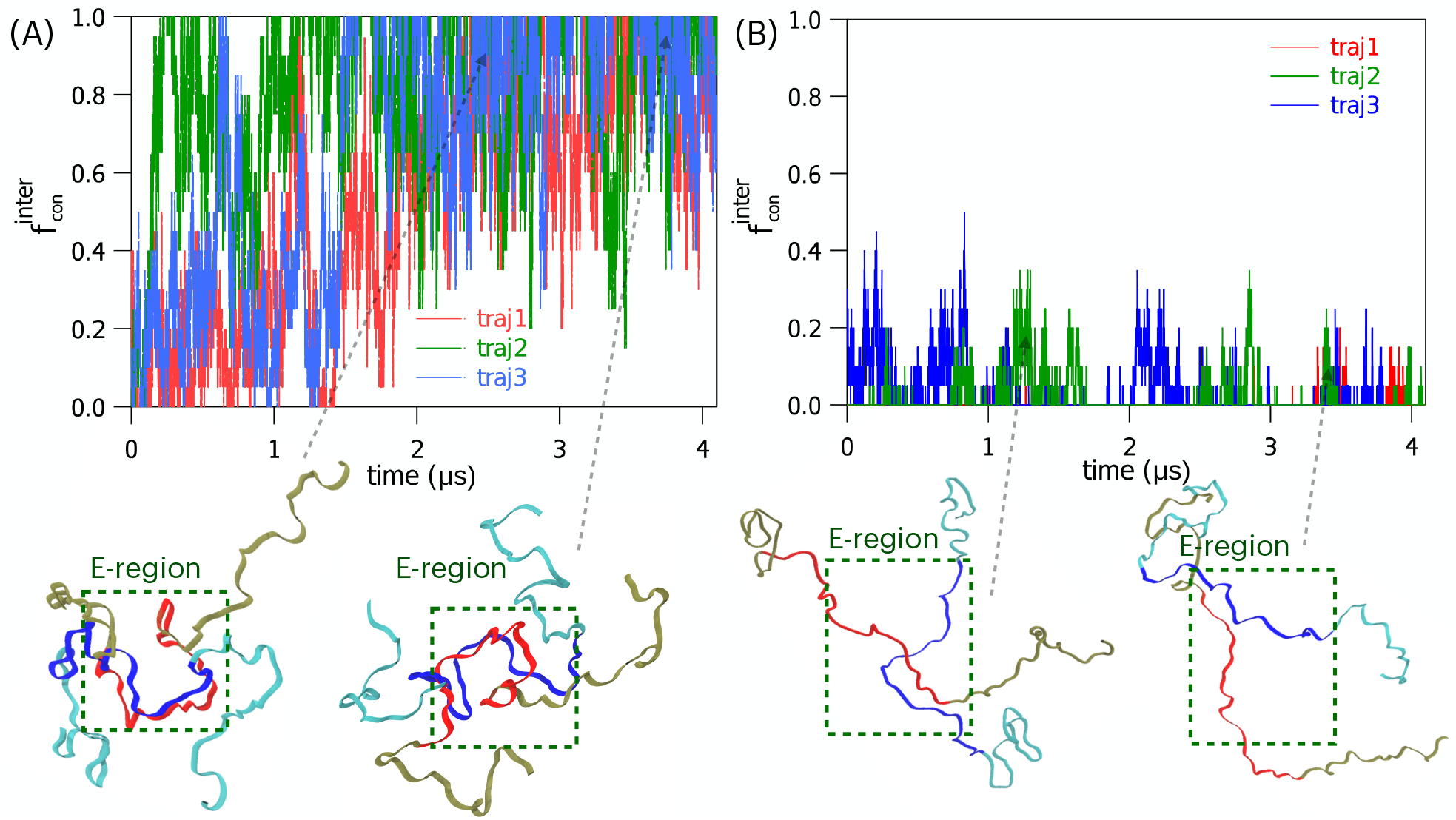
Dimer simulations were initiated with *N*^*∗*^ and non-*N*^*∗*^ states at acidic pH. (A) Fraction of interchain contact 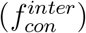 as a function of time is shown for three independent trajectories starting from *N*^*∗*^ states in red, green, and blue lines. The *N*^*∗*^ states reaches 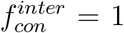 forming dimer within 1.5 *μ*s. The representative dimer conformations are shown in ribbon representation where the E-regions of chain-1 and chain-2 are given in blue and red, respectively and highlighted using a green rectangle. (B)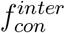 as a function of time is shown for three independent simulation trajectories starting from non-*N*^*∗*^ states in red, green, and blue lines. Trajectories initiated with non-*N*^*∗*^ states do not approach 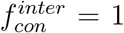 and mostly remain in the monomer state within the simulation timescale.

## Plausible mechanism of fibril formation at low pH

Based on our observations, we provide a plausible mechanism for proT*α* fibril formation at acidic pH (Figure 6). ProT*α* exists as an extended state at neutral pH due to the electrostatic repulsion between the negatively charged residues present mainly in the E-region of the IDP. With decreasing pH, there is compaction in the proT*α* size due to the neutralization of negatively charged residues. Low pH induces compaction in the protein conformations and enhances *β*-sheet content in the E-region. The conformations with high *β*-sheet content act as aggregation-prone *N*^*∗*^ conformations, which initiate oligomer nucleation that leads to fibril formation in the later stages of the aggregation process. From these results, we can further hypothesize that the E-region can form the fibril core in the aggregates, which can be verified using experiments.

## Conclusions

The pH-induced aggregation of IDPs is linked to multiple neurodegenerative diseases. Therefore, it is essential to understand how pH influences the conformational sampling of the IDPs and identify the aggregation-prone regions in the IDPs which are critical to the formation of amyloid fibrils. We answered the following questions related to pH-dependent IDP aggregation: (1) which region of proT*α* is affected by the pH change, and how does it influence the proT*α* conformational ensemble? Using a coarse-grained IDP model and molecular dynamics simulations, we showed that the intermediate region of the IDP proT*α* rich in glutamic acid residues becomes the aggregation-prone region on pH change from neutral to acidic. Due to the protonation of the glutamic acid residues at low pH, the propensity of *β*-sheet formation in the proT*α* intermediate region increases, and this region could be critical in the nucleation for the aggregate formation. We showed that the conformations with high *β*-sheet content form dimers readily and support the hypothesis that the intermediate region of proT*α* is critical to aggregation. The regions in the IDP that can have significant compaction on pH change are the patches of IDP that undergo a substantial change in NCPR. (2) Is there a relation between pH-induced compaction and aggregation? Our findings show that although the dominant fraction of the aggregation-prone *N*^*∗*^ conformations at low pH are compact, a minor population of *N*^*∗*^ conformations is extended, suggesting that the *N*^*∗*^ ensemble is heterogeneous.

**Figure 6:**
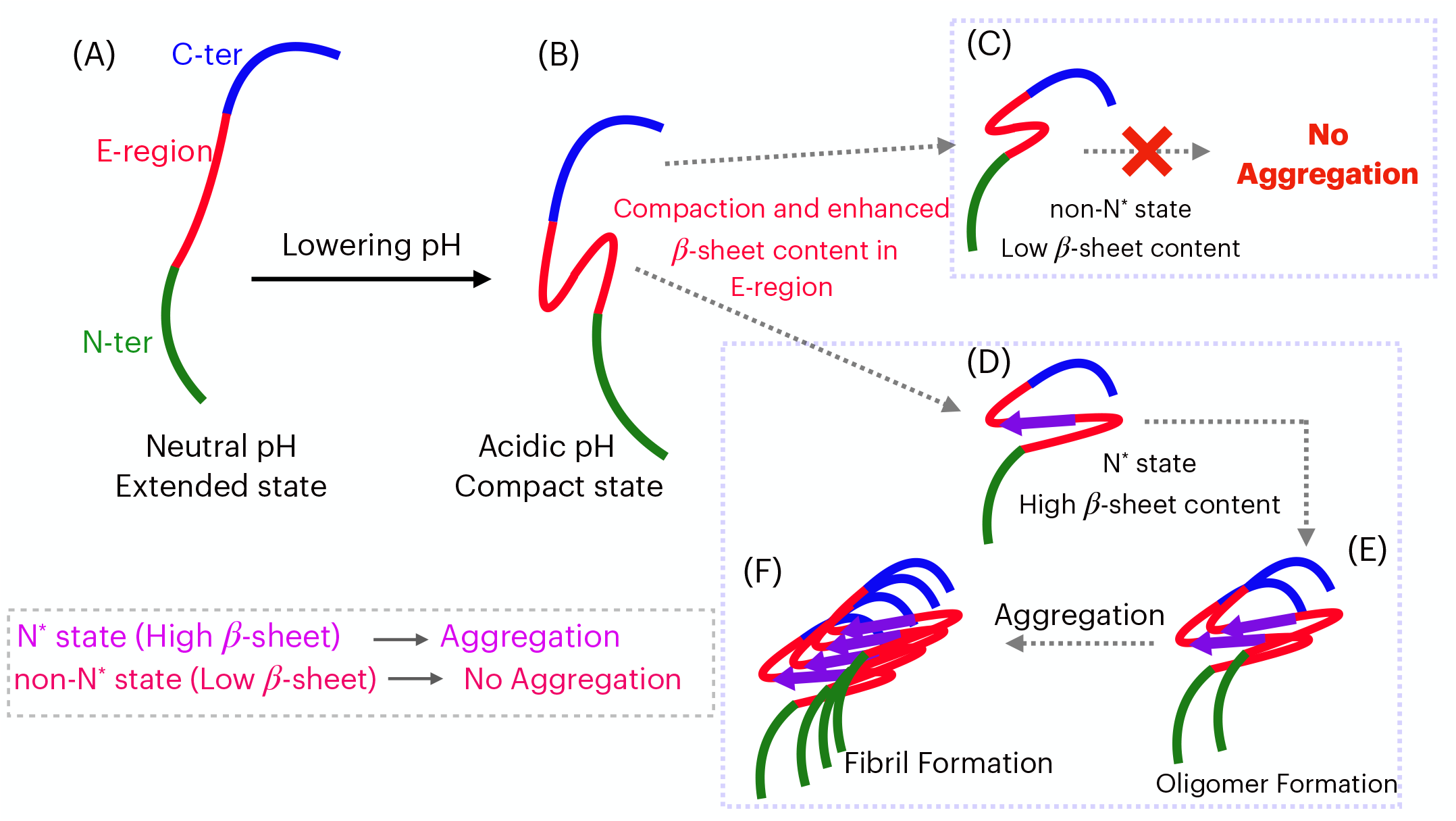
A plausible mechanism of proT*α* aggregation at acidic pH. (A) ProT*α* exists as an extended state at neutral pH due to the electrostatic repulsion. Three segments of proT*α* are shown in blue (C-ter), red (E-region), and green (N-ter). (B) pH change from neutral to acidic induces compaction and enhances *β*-sheet content specifically in the E-region. (C) Non-*N*^*∗*^ states with low *β*-sheet content do not aggregate. (D) The *N*^*∗*^ states with high *β*-sheet content can initiate the aggregation process. (E) The *N*^*∗*^ states form oligomers, where the E-region can act as a fibril core. (F) The oligomers mature into regular fibrils.

A similar pH-induced aggregation is also observed in other IDPs such as *α*-synuclein^67^ and tau.^68^ In *α*-synuclein, the NAC region of the IDP compacts upon pH change from neutral to acidic, and this region forms the fibril core. Combining these results, we can generalize that in the class of IDPs, which aggregate on pH change from neutral to acidic, the region in the IDP with the maximum compaction on pH change is critical to aggregation and forms the fibril core. These insights show that a plausible way to prevent the aggregation of such IDPs is to design new molecules that can bind to the IDP regions where maximum compaction is observed at low pH when the dominant acid residues are protonated.

## Materials and Methods

### Self Organized Polymer Model for Intrinsically Disordered Proteins (SOP-IDP)

We used a coarse-grained SOP-IDP^14,36^ model to study the pH effect on proT*α*. In this model, each residue is represented by two beads, one for the backbone atoms and another for the side chain atoms. The energy function of an IDP with bead coordinates, {**r**} for a protein conformation consists of three terms - bonded interaction (*E*_*B*_), local nonbonded 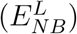 interaction nonlocal nonbonded interaction 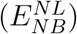 and electrostatic interaction (E_ele_)

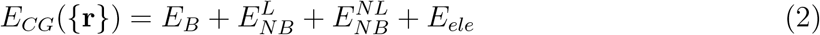

The bonded interaction is modeled using finitely extensible non-linear elastic (FENE) potential. The local nonbonded interaction is a purely repulsive potential to avoid overlapping among nonbonded beads. The nonlocal nonbonded interaction is modeled using the Lenard-Jones (LJ) potential. This potential exists between backbone-backbone, backbone-side chain, and side chain-side chain beads, which are nonbonded and separated by at least 2 residues. Electrostatic interaction is modeled using the Debye-Huckel screening potential. At neutral pH (pH ≈ 7.5), the charge for positively charged Lys and Arg residues is +1, and negatively charged Asp and Glu residues are is -1. At acidic pH (pH ≈ 2.5), negatively charged residues are neutralized, and the charge of Asp and Glu is 0. A detailed description of each term in the energy function (Eq 2) and various energy parameters are given in the supporting information (SI).

### Simulation Details and Data Analysis

We carried out low friction Langevin dynamics^69^ simulation at neutral and acidic pH by changing the charge of negatively charged residues at [*salt*] = 150 mM and temperature *T* = 298 K. The dimerization simulations are performed using Brownian dynamics simulations at the same conditions. The simulation details are given in the SI.

The IDP ensemble is characterized with the small–angle X–ray scattering (SAXS) intensity profiles (*I*(*q*)). The SAXS intensity is calculated using the equation

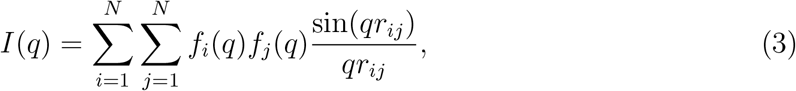

where *q* is the wave vector, *r*_*ij*_ is the distance between the beads *i* and *j, N* is the number of beads in the IDP and *f*_*i*_(*q*) is the form factor of bead *i* and their values are taken from ref.^70^

The normalized structure factor, ^51^ *S*(*q*), is calculated using the equation

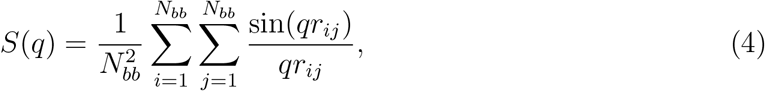

where *N*_*bb*_ is the number of backbone beads in the IDP, and *r*_*ij*_ is the distance between the backbone beads of residues *i* and *j*.

## Acknowledgement

GR acknowledges funding from the National Supercomputing Mission (MeitY/R&D/HPC/2(1)/2014). LB acknowledges research fellowship from the prime minister’s research fellows (PMRF) scheme. We acknowledge National Supercomputing Mission (NSM) for providing computing resources of “PARAM Brahma” at IISER Pune, which is implemented by C-DAC and supported by the Ministry of Electronics and Information Technology (MeitY) and Department of Science and Technology (DST), Government of India.

## Supporting Information

Supporting Information Available

