## Supporting Information for "pH Induced Switch in the Conformational Ensemble of an Intrinsically Disordered Protein Prothymosin-*α* and Its Implications to Amyloid Fibril Formation"

### Supporting Information for "pH Induced Switch in the Conformational Ensemble of an Intrinsically Disordered Protein Prothymosin-$\alpha$ and Its Implications to Amyloid Fibril Formation"

Lipika Baidya and Govardhan Reddy*

*Solid State and Structural Chemistry Unit, Indian Institute of Science, Bengaluru, Karnataka, India 560012*

#### Self Organized Polymer Model for Intrinsically Disordered Proteins (SOP-IDP)

We performed the simulations using a self-organized polymer model for intrinsically disordered proteins (SOP-IDP) described elsewhere.[1,2] In the SOP-IDP model, each residue is represented as two beads - one bead for the backbone atoms and the other bead for side chain atoms. The backbone and side-chain beads are situated at the $C_\alpha$ position and the center of mass of side chain atoms, respectively. To get the initial coarse-grained coordinates ($\{\mathbf{r}\}$) for the polypeptides, we generated an elongated chain with the polypeptide backbone beads pointed along the $Z$-direction.

The energy function ($E_{CG}(\{\mathbf{r}\},0)$) of the SOP-IDP model is the sum of bonded ($E_B$) and non-bonded ($E_{NB}$) and electrostatic ($E_{ele}$) interactions. The non-bonded energy consists of local ($E_{NB}^{L}$) and non-local ($E_{NB}^{NL}$) interactions. The Hamiltonian of the SOP-IDP model is

$$E_{CG}(\{\mathbf{r}\},0) = E_B + E_{NB}^{L} + E_{NB}^{NL} + E_{ele} \tag{S1}$$

The interaction between two bonded beads ($E_B$) is modeled using the finite extensible non-linear elastic (FENE) potential,

$$E_B = -\sum_{i=1}^{N_B} \frac{k}{2} R_0^2 \log\left(1 - \frac{(r_i - r_i^0)^2}{R_0^2}\right), \tag{S2}$$

where $N_B$ is the total number of bonds in the SOP-IDP model. $r_i$ is the instantaneous bond distance between the $i^{th}$ pair of bonded beads and $r_i^0$ is the corresponding equilibrium bond distance, and $R_0$ is the maximum bond extension/compression. The values of $r_i^0$ are set to the sum of van der Waals radii. The values of $k$ and $R_0$ are given in Table S1.

The two beads, which are not connected by a covalent bond and are separated by less than two residues along the polypeptide chain, interact with each other through a nonbonded local potential, which is purely repulsive to account for the excluded volume interactions to prevent unphysical overlap between the two nonbonded beads and is given by

$$E_{NB}^{L} = \sum_{i=1}^{N_l} \epsilon_l \left(\frac{\sigma_i}{r_i}\right)^6. \tag{S3}$$

Here $N_l$ is the number of local nonbonded interaction pairs, $\sigma_i$ is the sum of the van der Waals (vdW) radii of $i^{th}$ pair of nonbonded beads, and $\epsilon_l$ is the strength of repulsive interaction. The value of $\epsilon_l$ is given in Table S1, and values of vdW radii for each amino acid residue are listed in Table S2.

The beads, separated by more than two residues, interact through a nonbonded nonlocal interaction modeled using the Lenard-Jones type potential and is given by

$$\begin{aligned} E_{NB}^{NL} = & \sum_{i=1}^{N_{bb}} \epsilon_{bb} \left[\left(\frac{\sigma^{bb}}{r_i}\right)^{12} - 2\left(\frac{\sigma^{bb}}{r_i}\right)^{6}\right] \\ & + \sum_{i=1}^{N_{bs}} \epsilon_{bs} \left[\left(\frac{\sigma_i^{bs}}{r_i}\right)^{12} - 2\left(\frac{\sigma_i^{bs}}{r_i}\right)^{6}\right] \\ & + \sum_{i=1}^{N_{ss}} \epsilon_{ss} \mid 0.7 - \epsilon_i \mid \left[\left(\frac{\sigma_i^{ss}}{r_i}\right)^{12} - 2\left(\frac{\sigma_i^{ss}}{r_i}\right)^{6}\right]. \end{aligned} \tag{S4}$$

The first, second and third terms of Eq. S4 correspond to the backbone-backbone, backbone-side chain and side chain-side chain interactions energies, respectively. $N_{bb}$, $N_{bs}$ and $N_{ss}$ denote the number of interaction pairs present between backbone-backbone, backbone-side chain and side chain-side chain beads, respectively. $r_i$ is the distance between $i^{th}$ pair of beads. $\sigma^{bb}$ is the diameter of the backbone bead, which is 3.8 Å. $\sigma_i^{bs}$ and $\sigma_i^{ss}$ are the sum of bead radii for the $i^{th}$ pair of backbone-side chain and side chain-side chain beads, respectively. $\sigma_i^{bs}$ and $\sigma_i^{ss}$ are computed using the bead radii listed in Table S2. $\epsilon_i^{bb}$, $\epsilon_i^{bs}$ and $\epsilon_i^{ss}$ are the strength of backbone - backbone, backbone - side chain and side chain - side chain interactions, respectively. We used Betancourt - Thirumalai statistical potential[3] for $\epsilon_i$, which was initially proposed for the side chain - side chain interactions of globular proteins.

If an IDP contains charged residues, the beads corresponding to the side chains of the charged residues interact through a screened Coulomb potential given by

$$E_{ele} = \sum_{i=1}^{N_c-1} \sum_{j>i}^{N_c} \frac{q_i q_j \exp(-\kappa r_{ij})}{\epsilon r_{ij}}, \tag{S5}$$

where $N_c$ is the total number of charged residues present in the IDP, $r_{ij}$ is the distance between charged beads $i$ and $j$, $q_i$ and $q_j$ are the point charges measured in units of electron charge placed on the centers of the side chain beads $i$ and $j$, respectively. At neutral pH ($\approx$ 7.5), $q_i$ is +1 for positively charged Lys and Arg residues and -1 for negatively charged Asp and Glu residues. At acidic pH ($\approx$ 2.5), negatively charged residues neutralized and $q_i$ for Asp and Glu are considered as 0. The inverse Debye length, $\kappa$, is computed for a monovalent salt of concentration 150 mM. We have used the dielectric constant of the medium $\epsilon = 78.0$ $\epsilon_0$, where $\epsilon_0$ (= 1.0) is the permittivity of vacuum.

### Simulation Details and Data Analysis

#### Langevin Dynamics

We carried out low friction Langevin dynamics[4] simulation at temperature, $T$ = 300 K to compute the average thermodynamic properties of IDPs. The equation of motion in Langevin dynamics is given by,

$$m\ddot{\vec{r}} = -\zeta\dot{\vec{r}} + \vec{F}_C + \vec{\Gamma}, \tag{S6}$$

where $m$ is the mass of protein beads, $\zeta$ is the friction coefficient of the solvent medium, $\vec{F}_C$ is the deterministic force given by $-\frac{\partial E_{CG}(\{\mathbf{r}\},0)}{\partial \vec{r_i}}$, and $\vec{\Gamma}$ is the random force with Gaussian noise characterized by $\left\langle \vec{\Gamma}(t) \cdot \vec{\Gamma}(t+nh) \right\rangle = \frac{2\zeta k_B T}{h}\delta_{0,n}$ where $n = 0, 1, \cdots$, $\delta_{0,n}$ is the Kronecker delta function and $k_{\mathrm{B}}$ is the Boltzmann constant. We integrated Eq. S6 using the velocity Verlet algorithm. We used $\zeta = 0.05\ m/\tau_{\mathrm{LD}}$ and an integration timestep, $h = 0.005$ $\tau_{\mathrm{LD}}$, where, $\tau_{\mathrm{LD}}$ $\left(= \sqrt{\frac{m_0 a_0^2}{\epsilon_h}}\right)$ is the unit of time used to advance the Langevin dynamics simulations. The average mass of each bead ($m_0$), characteristic unit of length ($a_0$) and energy ($\epsilon_h$) are taken as 1.8 $\times 10^{-22}$ g, 1 Å and 1 kcal/mol, respectively. The value of $\tau_{\rm LD}$ in real time unit is $\approx$ 1.3 ps. We have at least $10^5$ snapshots for each system to calculate the thermodynamic properties.

#### Brownian Dynamics

We performed the dimer simulations using Brownian dynamics. We initiated the simulation by placing two randomly chosen monomer conformations at $\langle R_g \rangle$ ($\approx$ 30 Å) distance. To maintain the critical concentration for dimer formation, we added a harmonic potential between the centre of mass (COM) of two chains with a spring constant $k_{spring} = 2.0$ kcal/mol/Å$^2$ to maintain the COM distance between the two chains less than $\langle R_g \rangle$. The harmonic potential is active if the COM distance between the two chains is greater than their ensemble average $\langle R_g \rangle$. The equation of motion is integrated using Ermak-McCammon algorithm given by

$$\vec{r}_i(t+h) = \vec{r}_i(t) + \frac{h}{\zeta_H}\vec{F}_C + \vec{\Gamma}, \tag{S7}$$

where $\vec{\Gamma}$ is a random displacement of gaussian distribution with mean zero and variance $\langle \Gamma(h)^2 \rangle = \frac{2k_BTh}{\zeta_H}$. The friction coefficient $\zeta_H = 21.9\ m/\tau_{BD}$ and $h = 0.001\ \tau_{BD}$. The real simulation timescale $\tau_{BD} \sim \frac{\zeta_H a^2}{k_BT} = \frac{(\zeta_H \tau_{LD}/m)\epsilon}{k_BT}\tau_{LD}$.

#### Data Analysis

The simulated small-angle X-ray scattering (SAXS) intensity ($I(q)$) profiles for the IDPs are calculated using the expression

$$I(q) = \sum_{i=1}^{N}\sum_{j=1}^{N} f_i(q) f_j(q) \frac{\sin(qr_{ij})}{qr_{ij}}, \tag{S8}$$

where $N$ is the total number of beads, $q$ is the scattered wave vector, $f_i(q)$ is the form factor of bead $i$, and $r_{ij}$ is the distance between the beads $i$ and $j$. The values of $f_i(q)$'s are taken from ref.[5]

The radius of gyration, $R_g$, of the IDPs is given by

$$R_g = \left( \frac{1}{2N^2} \sum_{i,j} \vec{r}_{ij}^{\,2} \right)^{1/2}, \tag{S9}$$

where $\vec{r}_{ij}$ is the vector joining beads $i$ and $j$ and $N$ is the number of beads in the IDP. The normalized structure factor,[6] $S(q)$, is computed using the equation

$$S(q) = \frac{1}{N_{bb}^2} \sum_{i=1}^{N_{bb}} \sum_{j=1}^{N_{bb}} \frac{\sin(qr_{ij})}{qr_{ij}}, \tag{S10}$$

where $N_{bb}$ is the number of backbone beads in the IDP, and $r_{ij}$ is the distance between the backbone beads of residues $i$ and $j$.

(5) Tong, S., D. Yang; Lu, L. Accurate optimization of amino acid form factors for com-

puting small angle X-ray scattering intensity of atomistic protein structures. *J. Appl. Crystallogr.* **2016**, *49*, 1148–1161.

(6) Rubinstein, M.; Colby, R. *Polymer Physics*; Oxford University Press Oxford: New York, 2003.

Table S1: Parameters used in SOP-IDP Model

| Parameter | Value |
|---|---|
| $k$ | 20.0 kcal/(mol.Å$^2$) |
| $R_0$ | 2.0 Å |
| $\epsilon_l$ | 1.0 kcal/mol |
| $\epsilon_{bb}$ | 0.12 kcal/mol |
| $\epsilon_{bs}$ | 0.24 kcal/mol |
| $\epsilon_{ss}$ | 0.18 kcal/mol |

Table S2: Backbone and side chain bead radii of amino acid residues

| Bead | vdW radius (Å) |
|---|---|
| backbone | 1.90 |
| Gly | 0.5 |
| Ala | 2.52 |
| Val | 2.93 |
| Leu | 3.09 |
| Ile | 3.09 |
| Met | 3.09 |
| Phe | 3.18 |
| Pro | 2.78 |
| Ser | 2.59 |
| Thr | 2.81 |
| Asn | 2.84 |
| Gln | 3.01 |
| Tyr | 3.23 |
| Trp | 3.39 |
| Asp | 2.79 |
| Glu | 2.96 |
| Hse | 3.04 |
| Hsd | 3.04 |
| Lys | 3.18 |
| Arg | 3.28 |
| Cys | 2.74 |

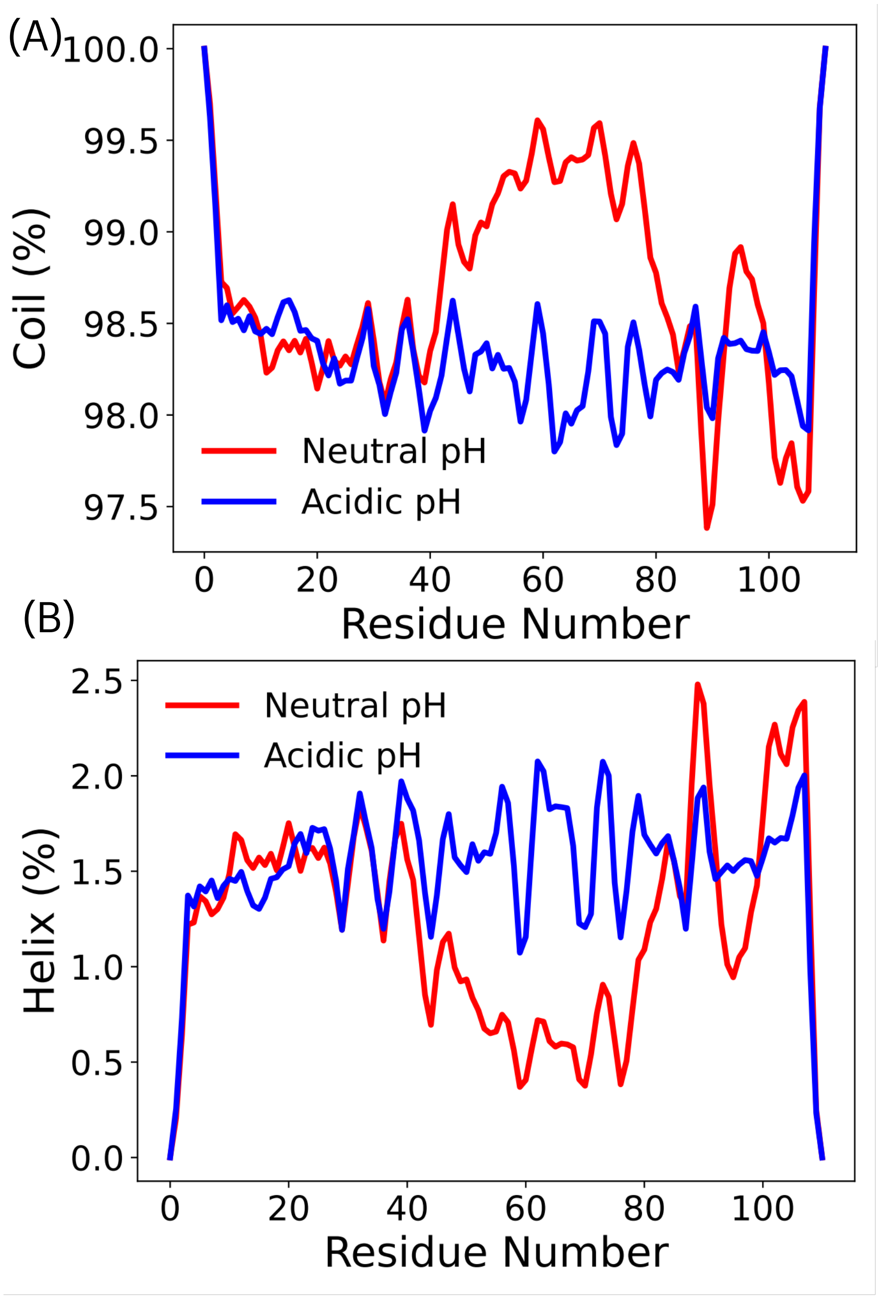


Figure S1: Percentage of (A) coil and (B) helix content as a function of residue number at neutral (red) and acidic (blue) pH. The helix content increases whereas coil content decreases at acidic pH in the E-region (E41 - D80).

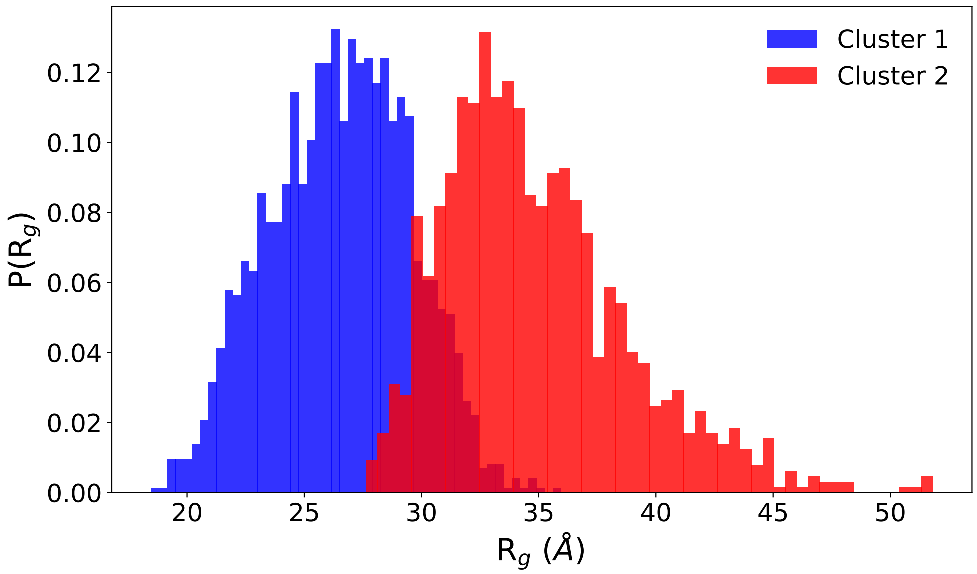


Figure S2: Probability distribution of $R_g$, $P(R_g)$, of the two $N^*$ clusters obtained from t-SNE dimensionality reduction technique. Cluster 1 and 2 contain compact and extended conformations, respectively.

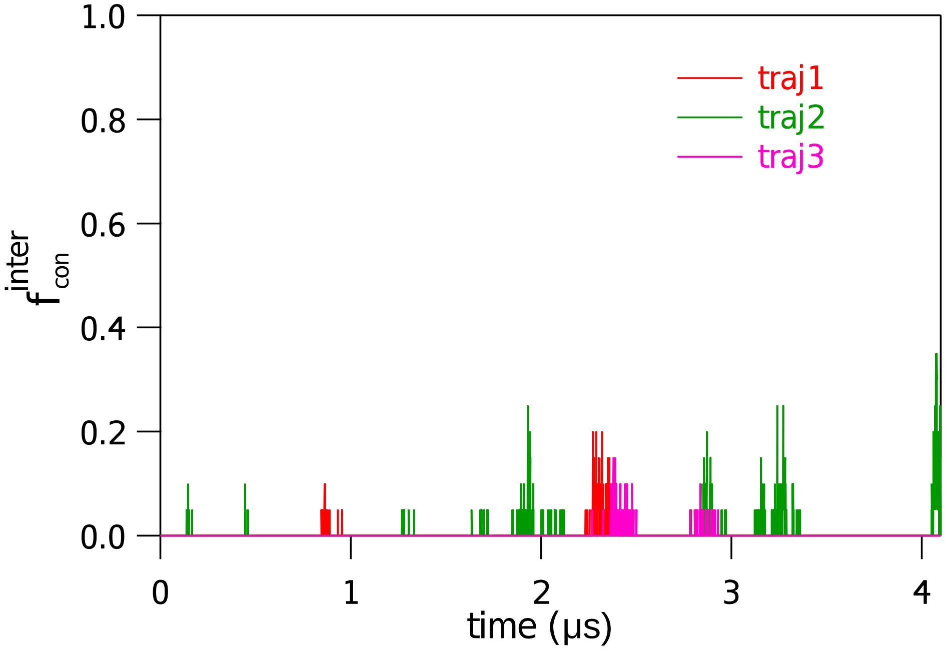


Figure S3: Fraction of interchain contact ($f_{con}^{inter}$) as a function of simulation time at neutral pH ($\approx$ 7.5) for three independent trajectories shown in red, green and magenta line. There is no signature of dimer formation as $f_{con}^{inter}$ never reaches 1 and hops between 0 to 0.4.
